# The Development, but not Expression, of Alcohol Front-loading in C57BL/6J Mice Maintained on LabDiet 5001 is Abolished by Maintenance on Teklad 2920x Rodent Diet

**DOI:** 10.1101/2022.02.23.481358

**Authors:** Nicole M. Maphis, Tameryn Radcliff Huffman, David N. Linsenbardt

## Abstract

Excessive alcohol (ethanol) consumption, such as binge-drinking, is extremely commonplace and represents a major health concern. Through modeling excessive drinking in rodents, we are beginning to uncover the neurobiological and neurobehavioral causes and consequences of this pattern of ethanol intake. One important factor for modeling binge drinking in mice is that subjects reliably drink to blood ethanol concentrations (BECs) of 80 mg/dl or higher. Drinking-in-the-dark (DID) is a commonly used mouse model of binge drinking, and we have shown these methods reliably result in robust ethanol front-loading and binge-level BECs in C57BL/6J (B6) mice as well as other ethanol-preferring mouse strains/lines. However, establishing the DID model in a new vivarium space forced us to consider the use of rodent diet formulations we had not previously used. The current set of experiments were designed to investigate the role of two standard rodent diet formulations on binge drinking and the development of ethanol front-loading using DID. We found that BECs in animals maintained on LabDiet 5001 (LD01) were double those found in mice maintained on Teklad 2920x (TL20). Interestingly, this effect was paralleled by differences in the degree of front-loading, such that LD01-fed mice consumed approximately twice as much ethanol in the first 15 minutes of the 2-hour DID sessions compared to TL20-fed mice. Surprisingly however, mice that developed front-loading during maintenance on the LD01 diet continued to display front-loading behavior after being switched to the TL20 diet. These data emphasize the importance of choosing and reporting diet formulations when conducting voluntary drinking studies and support the need for further investigation into the mechanisms behind diet-induced differences in binge drinking, particularly front-loading.

## Introduction

A large percentage of mortality and disease, globally, can be attributed to alcohol (ethanol) use. In the US alone, excessive ethanol drinking leads to 95,000 deaths and more than $250 billion in financial costs each year (numbers from 2010; CDC). Binge drinking, defined as a pattern of drinking that leads to a blood ethanol concentration (BAC) at or over 0.08 g/dL (or 80 mg/dL; NIAAA/NIH), is a particularly detrimental form of ethanol use. Unfortunately, binge drinking is also extremely commonplace, and there are signs it may be increasing. For example, a survey conducted during lockdowns associated with the Covid-19 pandemic observed that not only did rates of ethanol consumption increase significantly in both males and females, but episodes of binge drinking also increased (Pollard et al., 2020), especially amongst previous binge drinkers (Weerakoon et al., 2020). Another study corroborated these findings and found that these increases in general ethanol consumption and binge drinking during the Covid-19 pandemic also occurred broadly across all sociodemographic sub-groups (Barbosa et al., 2021). Thus, identifying factors that contribute to drinking too much ethanol, too quickly, is critical for developing strategies to improve public health.

Given ethical issues surrounding the imposition of binge ethanol exposure on human subjects genetically vulnerable to developing an alcohol use disorder (AUD), frequent unreliability of self-reporting in clinical studies, and limited access to human brain tissue, the use of reliable animal models are paramount to uncovering the neurobiological underpinnings of binge drinking. There are now many such rodent models, each of which has its own advantages (Belknap et al., 1993; Bell et al., 2006; Fuller, 1964; Gill et al., 1996; Grahame et al., 1999; Lim et al., 2012; McClearn et al., 1964; Rhodes et al., 2005). Certain strains of mice provided with ethanol access using one such model, known as ‘Drinking-in-the-Dark’ (or ‘DID’), reliably consume ethanol to binge-level BACs, display escalations in binge ethanol consumption and tolerance with increasing DID experience, as well as ethanol “front-loading”; consumption of a disproportionately large amount of ethanol shortly following ethanol availability (Ardinger et al., 2020; Linsenbardt et al., 2011; Linsenbardt and Boehm, 2015, 2014, 2012). While each of these phenotypes are frequently observed in binge drinking human clinical populations, we have developed a keen interest from our own studies over the last decade in front-loading, which we have previously hypothesized may reflect progressive increases in the motivation to experience the positive post-absorptive effects of ethanol (Ardinger et al., 2020; Linsenbardt and Boehm, 2014). Given that ongoing work in our lab as well as more recent clinical work suggests that ethanol front-loading may be a precursor to, or predictor of, ethanol use disorders (Carpenter et al., 2019), determining the mechanisms of ethanol front-loading, in particular, remains of great interest.

Among the many experimental variables capable of impacting motivation to voluntarily consume ethanol in excess, dietary components of rodent chow could be of significant importance. For example, greatly increasing the palatability of food has been shown to decrease ethanol consumption (Dole et al., 1985). However, more recently, seemingly subtle differences in rodent diets have been found to lead to profound differences in ethanol consumption (Marshall et al., 2015)(Quadir et al., 2020), including binge drinking using DID (Marshall et al., 2015). Thus, assessing ethanol consumption using DID in subjects maintained on rodent diets most frequently used in our animal research facility was a logical first step in setting up our research program at a new institution.

## Materials and Methods

### Animals

C57BL/6J mice (JAX Stock #000664) mice were purchased from The Jackson Laboratory (JAX) at 8 weeks of age and allowed to acclimate to a reverse light/dark cycle (12 hours off/on) and single-housing in standard ‘Allentown’ shoebox cages for at least two weeks prior to experimentation. Animals had *ad lib* access to one of two rodent diets (detailed below) and water, except during the 2-hour DID protocol, when (some) mice had access to a 20% ethanol solution, instead of water. DID procedures were initiated when mice were at least 70 days of age. All procedures were approved by the University of New Mexico Animal Care and Use Committee and conformed to the Guidelines for the Care and Use of Mammals in Neuroscience and Behavioral Research (2003).

### Diets

Animals were maintained on one of 2 standard rodent diets; Teklad 2920x (TL20; Envigo^®^, Madison, Wisconsin) or LabDiet 5001 (LD01; LabDiet^®^, St. Louis, MO). These diets are very similar in proportions of protein, carbohydrate, and fat, but possess a diverse variety of additional ingredients making up these proportions (see Table 1, Supplementary Table 1, Supplementary Table 2, and Supplementary Figure 1).

**Table 1.** Main Ingredients in two standard and readily available rodent diets.

| Abbreviated Name | LD01 | TL20 |
| --- | --- | --- |
| Manufacturer/name | Purina®/LabDiet 5001 | Envigo®/Teklad 2920x |
| Pellet Form | Compressed | Extruded |
| Soy-protein Concentration, mg/kg | 300-500 | Soy-Protein Free |
| Energy Density (kcal/g) | 3.02 | 3.1 |
| % Calories Provided by: |  |  |
| Protein | 28.507 | 24 |
| Fat (ether extract) | 13.496 | 16 |
| Carbohydrates | 57.996 | 60 |
| Main Ingredients (listed in descending order) | Ground corn, dehulled soybean meal, dried beet pulp, fish meal, ground oats, brewers dried yeast | Ground wheat, ground corn, corn gluten meal, wheat middlings, soybean oil |

### Drinking-in-the-Dark (DID)

DID procedures as performed here have been previously described in detail (Linsenbardt et al., 2011; Linsenbardt and Boehm, 2014, 2012, 2009); briefly, 3 hours into the dark cycle, bottles containing tap water were replaced with specialized sippers (monitor fluid consumption in real-time) containing either 20% ethanol solution (in tap water) or tap water, for a period of 2 hours. At the conclusion of the 2-hour DID period, bottles containing tap water were returned. DID procedures were repeated daily for 15 consecutive days.

### Blood Collection and Processing

For the determination of blood ethanol concentrations (BECs), peri-orbital sinus blood was drawn immediately following the last day of DID (day 15 or day 30), or 1- or 2-hours following experimenter administered ethanol on day 16 for assessment of ethanol metabolism (see below). Blood samples were then centrifuged at 10,000 RPM for 10 minutes, and plasma was withdrawn and stored at −20°C until enzymatic assessment of BECs using an Analox Ethanol Analyzer (Analox Instruments, Lunenburg, MA).

### Ethanol Metabolism

A subset of mice were administered 2.0 g/kg ethanol (20% v/v) via intraperitoneal (i.p.) injection on day 16, 24 hours following day 15 fluid access. Immediately following i.p. injection of 20% ethanol, mice were returned to their home cages for 1 or 2 hours, and then blood samples were collected and processed as detailed above for determination of BECs.

## Experiments

Experiment 1 was the first experiment conducted in our new lab and was designed to test our newly acquired and assembled volumetric sipper hardware/software using 12 male and 12 female mice. All mice were maintained on TL20 diet throughout and were given 15 consecutive 2-hour DID ethanol sessions. Experiment 2 used 40 female mice, maintained on either TL20 or LD01 rodent diets, and half of which were assigned to either ethanol or water DID solutions. Mice were given 15 consecutive 2-hour DID sessions before potential ethanol metabolism differences were assessed on the 16^th^ day. Experiment 3 used 20 female mice, half of which received either TL20 or LD01 rodent diets. Following 15 consecutive 2-hour DID ethanol sessions, mice in this experiment were then provided the opposite rodent diet for a period of two weeks, before another round of 15 consecutive 2-hour DID ethanol sessions were conducted.

## Results

### Experment 1: Rodents maintained on Teklad 2920x (TL20) do not reliably binge-drink

Repeated measures ANOVA did not detect any statistically significant differences in ethanol consumption as a function of sex (F(1,22)=2.36; p = 0.1384) or day (F(14,308)=0.71; p = 0.3240; Figure 1A). Assessment of day 15 ethanol intake (student’s t-test, p=0.3083) and consequent BEC (student’s t-test, p=0.9957; Figure 1B) also failed to detect any sex differences. Furthermore, and most importantly, mean BECs were well below the 80 mg/dl binge threshold (average of males and females = 39.43 ± 9 mg/dl), justifying experiments 2 and 3, which were designed specifically to test the hypothesis that mice maintained on LD01 would consume significantly more ethanol versus mice maintained on TL20.

**Figure 1.**
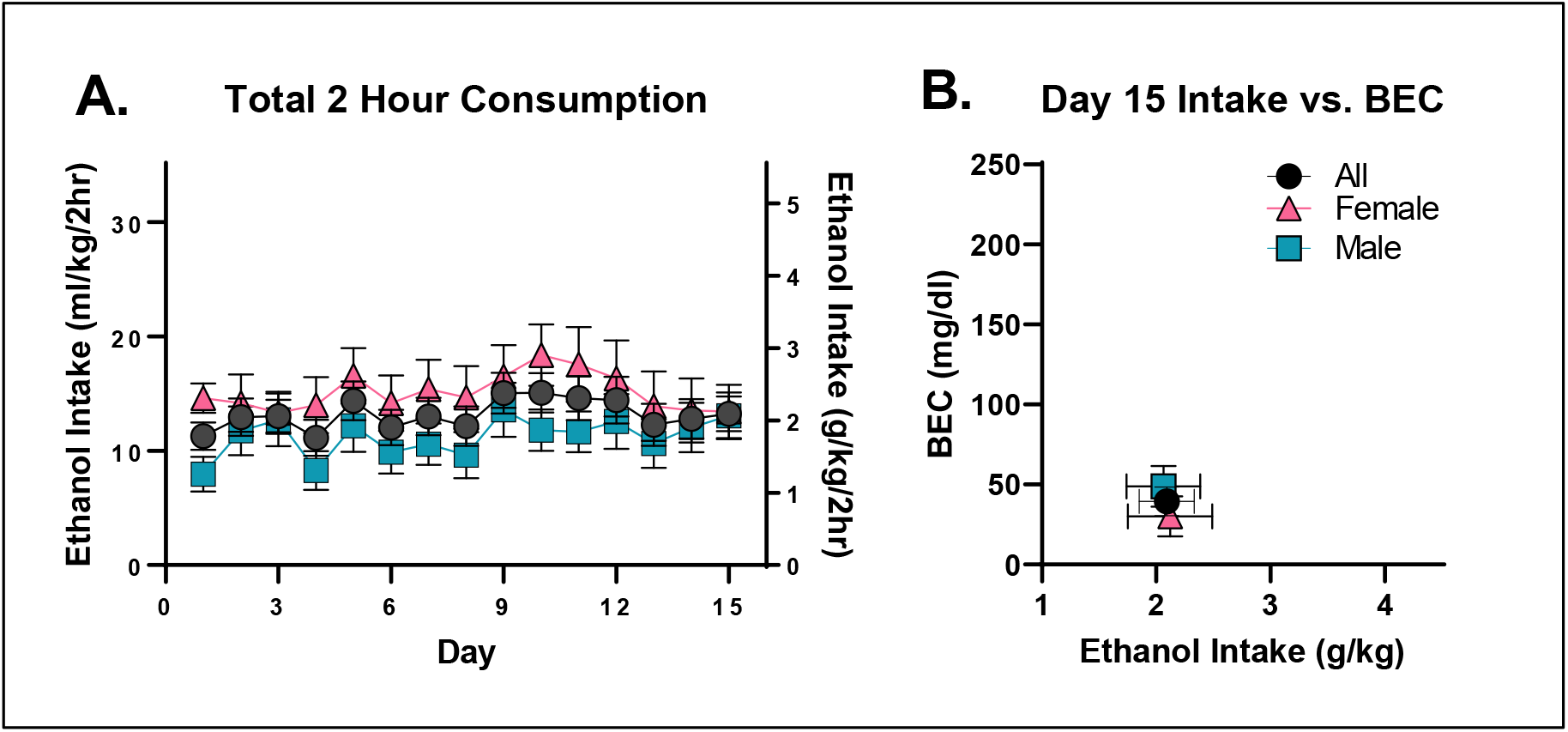
C57BL/6J mice do not reliably binge drink ethanol when maintained on TL20. **A**. Mean daily ethanol consumption over the two-hour consumption period as a function of sex. **B.** Day 15 BECs presented as a function of total ethanol consumed.

### Experiment 2. Increases in general fluid consumption in mice maintained on LD01

The influence of TL20 and LD01 rodent diets on ethanol consumption, water consumption, and BEC can be seen in Figure 2. Repeated-measures ANOVA of ethanol consumption identified a significant main effect of diet [F(1,18)=21.31; p=0.0002] and significant day*diet interaction [F(14,252)=2.21; p=0.0080] (Figure 2A). Similar results were observed in water consuming mice (Figure 2B) with repeated-measures ANOVA identifying a significant main effect of diet [F(1,18)=14.85; p=0.0012].

**Figure 2.**
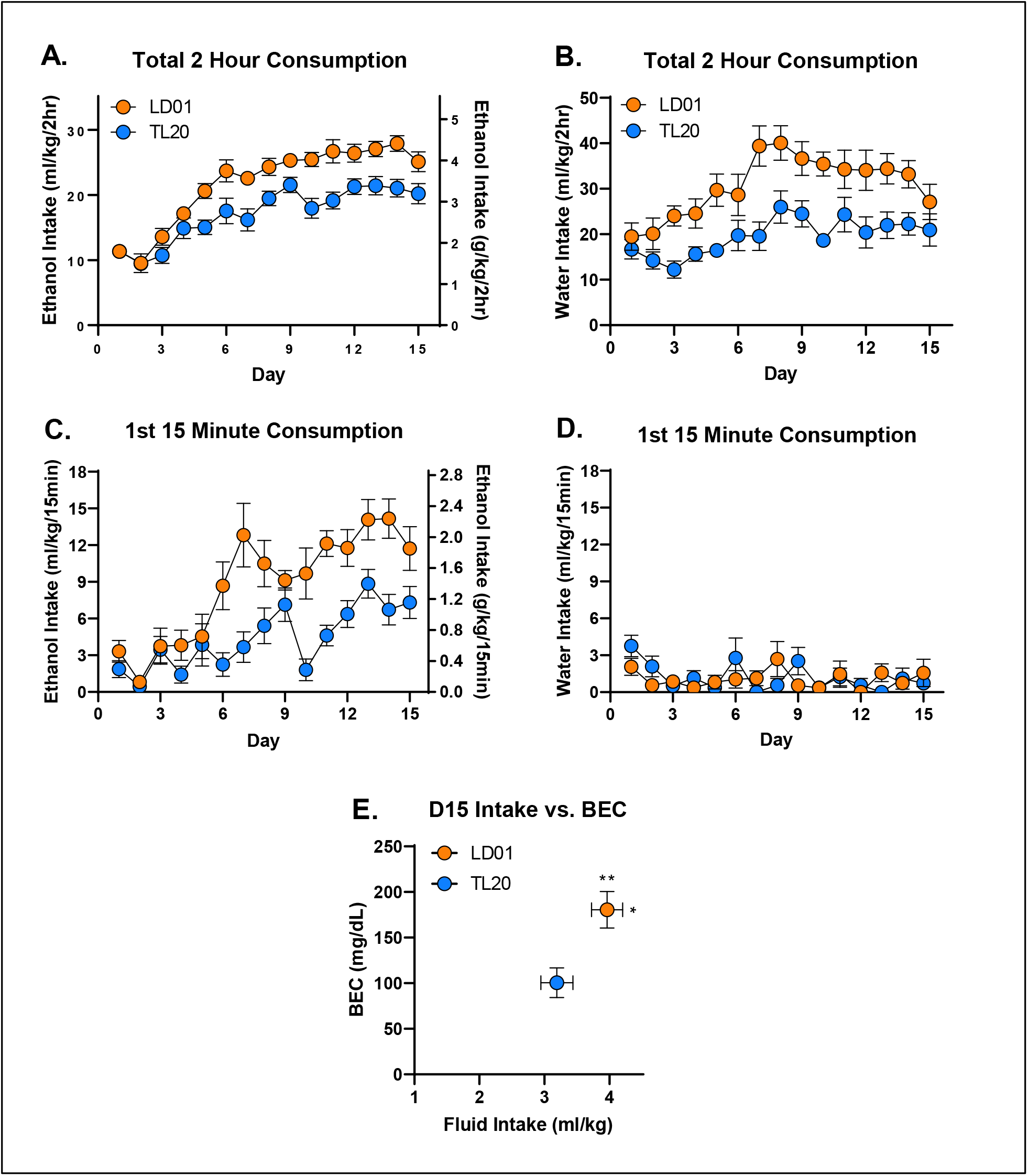
Fluid consumption is greater in mice fed LD01. Total 2-hour ethanol (**A**) and water (**B**) consumption in mice maintained on the LD01 was greater than in mice maintained on TL20. Mean ethanol consumption during the first 15 minutes of access was greater in mice fed LD01 (**C**), while mean water consumption during the first 15 minutes did not differ (**D**). Differences in ethanol consumption resulted in significant differences in BECs (**E**).

We next evaluated the impact of diet on drinking during the first 15 minutes of the session, as this is the conventional time period we have used to assess front-loading. Repeated measures ANOVA of ethanol consumption during the first 15 minutes identified a significant main effect of diet [F(1,18)=18.39; p=0.0004] and significant day*diet interaction [F(14,252)=2.94; p=0.0004], with LD01 consuming mice quickly developing greater front-loading compared to TL20 consuming mice (Figure 2C). An identical analysis for water consuming mice found no statistically significant effects (Figure 2D). Unsurprisingly, differences in ethanol intake (student’s t-test, p = 0.0374) were paralleled by differences in BEC (student’s t-test, p = 0.0062) (Figure 2E). Assessment of body weight found a significant main effect of diet on day 15 [F(1,36)=4.249; p=0.0466], but only when analyzed separately from Day 0, indicating that rodent diets, irrespective of fluid assignment, had a small but significant effect on weight gain over the approximately 2 week study, with slightly greater increases in mice consuming TL20 (Supplementary Figure 1A-C). Neither fluid assignment, nor diet consumed, significantly affected ethanol metabolism following an experimenter-administered ethanol challenge (Supplementary Figure 1D).

### Experiment 3: Rodents maintained on LD01 reliably consume more ethanol, display increased front-loading behavior, and achieve approximately two-fold higher BECs than animals maintained on TL20

The influence of TL20 and LD01 rodent diets on ethanol consumption, rate of ethanol consumption, and BEC can be seen in Figures 3–5. Repeated-measures ANOVA of all 4 weeks of ethanol consumption revealed a significant day*diet interaction [F(29,493)=11.94; p<.0001] which was driven by greater ethanol consumption in mice maintained on LD01 versus those maintained on TL20 during both the first 15 days of access (Figure 3B) and final 15 days of access (Figure 3D). This finding was further confirmed by repeated-measures ANOVAs for each 15 day block separately, which identified main effects of diet for each block [D1-15: F(1,18)=36.31; p<0.0001; D16-30: F(1,17)=5.009; p=0.0389] (Figures 3B, 3D). Differences in ethanol intake (student’s t-test, p = 0.0057) over the course of the first 15 days resulted in over two-fold difference in BECs (student’s t-test, p = 0.0007) (Figure 3C), with only animals in the LD01 group displaying binge-levels BEC as a group (i.e. >80 mg/dl). However, even though animals switched from TL20 to LD01 consumed significantly more ethanol than on the previous diet (Figure 3B, Figure 3D), these differences were much smaller in magnitude (student’s t-test, p = 0.3201) and did not lead to statistically significant differences in BECs (student’s t-test, p = 0.6333) (Figure 3E).

**Figure 3.**
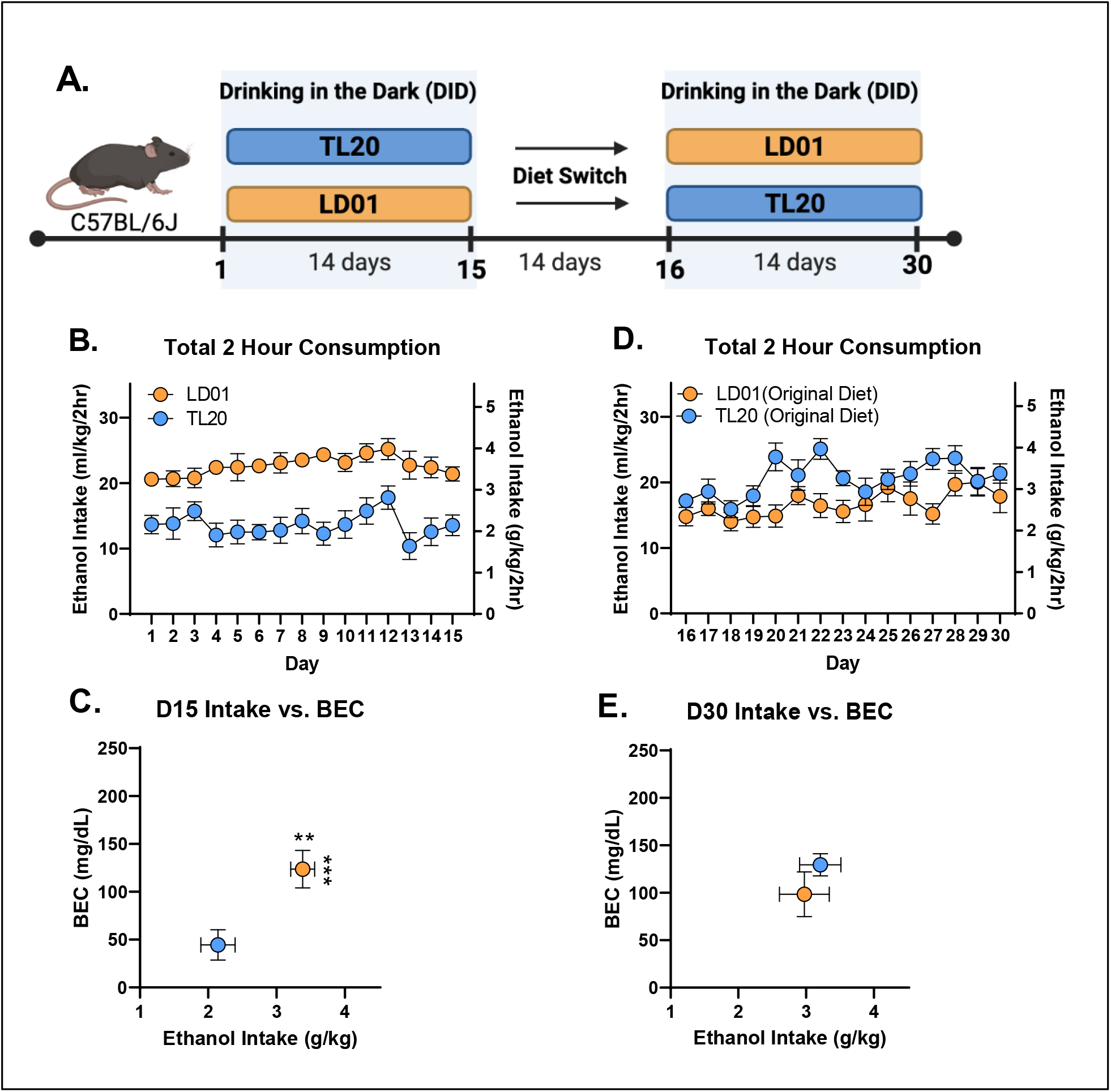
B6 mice maintained on LD01 reliably consume more ethanol and achieve approximately two-fold higher BECs versus animals maintained on TL20. **(A)** Animals were maintained on one of two diets (TL20 or LD01) during the first DID session, and then were switched to the opposite diet for a period of two weeks before undergoing a second DID session. **(B)** Mean daily 2-hour ethanol consumption over the first 15 days of access. **(C)** Day 15 BECs as a function of ethanol consumption **(D)** Mean daily 2-hour ethanol consumption over the second 15 day DID session. **(E)** Day 30 BECs as a function of ethanol consumption (** p< 0.01; ***p< 0.001).

**Figure 4.**
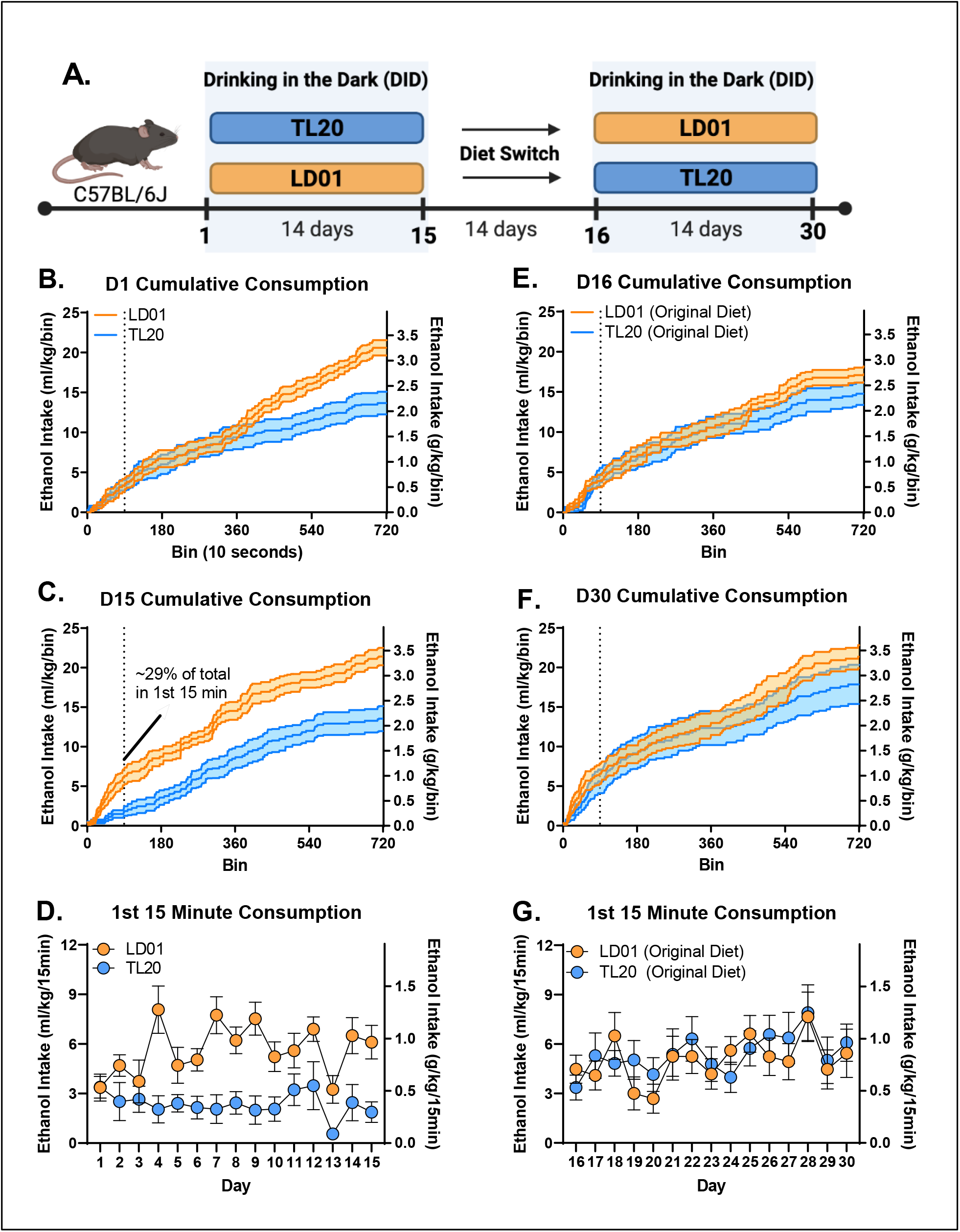
Differences in front-loading as a function of diet and previous experience with ethanol. **(A)** Animals were maintained on one of two diets (TL20 or LD01) during the first DID session (D1-D15), and then were switched to the opposite diet for a period of two weeks before undergoing a second DID session (D16-D30). Cumulative consumption over the 2 hour DID, dotted line indicates the first 15 min period within DID, on Day 1 (**B**) and D15 (**C**) demonstrates elevated ethanol front-loading in mice maintained on the LD01 diet compared to mice on the TL20 diet. (**D**) Differences in ethanol front-loading (first 15 minutes) emerged between LD01 and TL20 by day 4, and persisted through the first 15 days. (**E**) Mice were switched to the opposite diet for a period of 2 weeks before DID was conducted for a second block and cumulative consumption was measured over the 2 hour DID on Day 16 (**E**) and Day 30 (**F**). (**G**) Differences in ethanol front-loading (first 15 minutes) over block 2 (D16-D30) but overall ethanol consumption increased, especially in the mice switched from TL20 to LD01.

**Figure 5.**
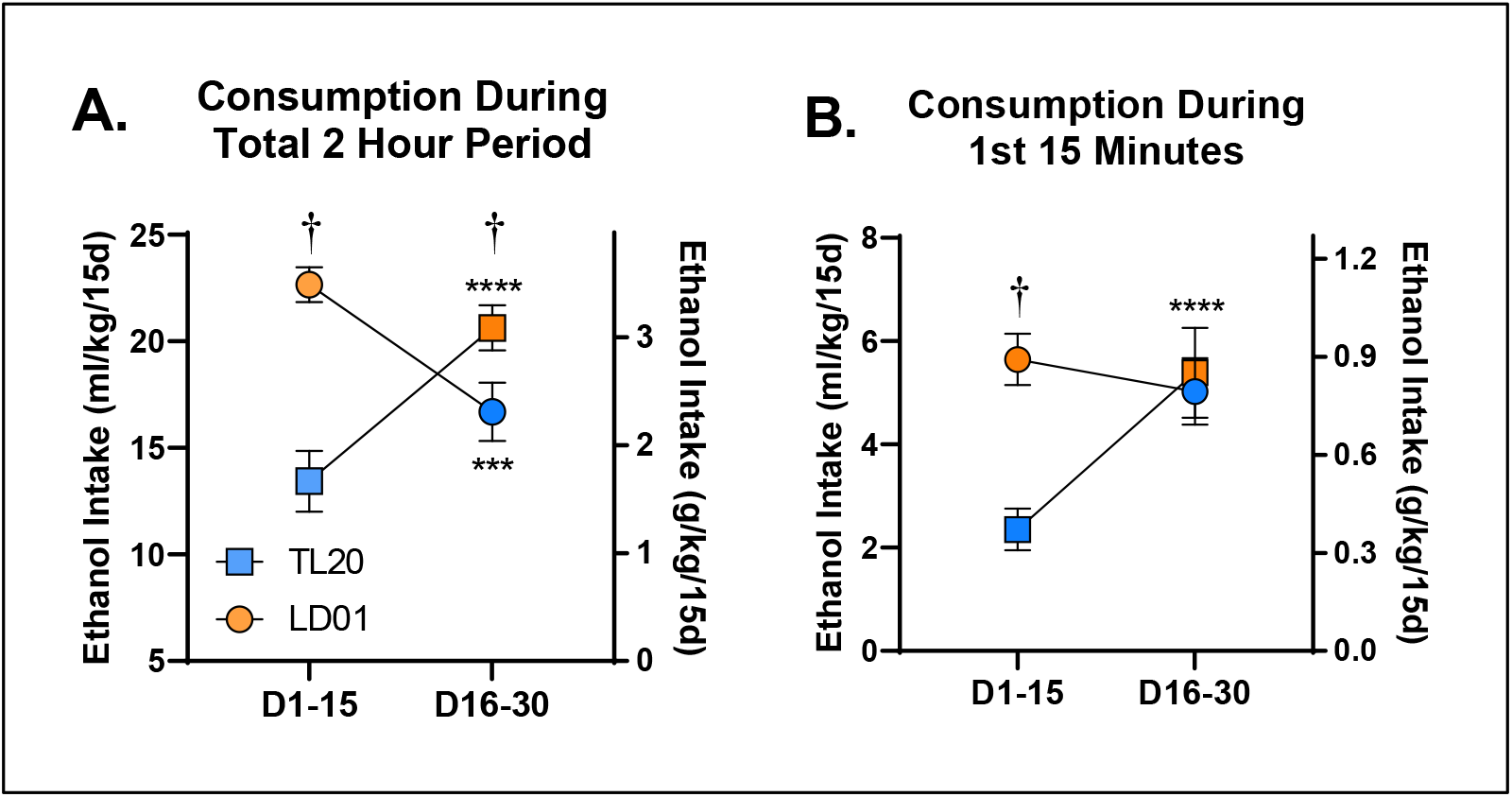
The development, but not expression, of ethanol front-loading in C57BL/6J mice maintained on Labdiet 5001 is abolished by maintenance on Teklad 2920x rodent diet. B6 mice switched from TL20 to LD01 diet display increases in general ethanol consumption and increases in front-loading compared with mice switched from LD01 to TL20 **(A)** Average 2-hour ethanol consumption during the first 2-week block versus the last 2-week block as a function of diet. Symbols (circle vs. square) indicate mouse cohort and colors (orange vs. blue) indicate diet assignment. †’s Indicate between-group differences [p < 0.05]; *’s Indicate within-group differences [***p<.001; ****p<.0001]. **(B)** Ethanol consumption during the first 15 minutes during the first 2-week block versus the last 2-week block as a function of diet. Symbols (circle vs. square) indicate mouse cohort and colors (orange vs. blue) indicate diet assignment. †’s Indicate between-group differences [p < 0.05]; *’s Indicate within-group differences [***p<.001; ****p<.0001].

To determine the extent to which differences in ethanol consumption might be driven by differences in ethanol drinking rate, i.e. front-loading, we next explored the time course of drinking throughout the daily 2-hour sessions (Figure 4A). Differences in mean ethanol intake between the two diet groups were initially mediated by a persistent drinking rate in the LD01 group (Figure 4B). However, differences in mean intake quickly shifted to being attributable to differences in front-loading; more specifically, lack of front-loading in the TL20 diet group (Figure 4C). Analysis of the first 15 minutes of ethanol consumption revealed a significant main effect of diet [F(1,18)=26.47; p<.0001], as well as a day*diet interaction [F(14,252)=2.022; p=0.0168], confirming the relative stability of differences in front-loading once they emerged (Figure 4D). We then explored the 2^nd^ 15-day block of drinking, and much to our surprise, did not observe any significant differences in ethanol front-loading (Figure 4E, Figure 4F), principally because front-loading in the group that had been established by mice previously maintained on LD01, continued front-loading once they were switched to TL20 diet, even by the last day of the 2^nd^ 15-day block (Day 30, Figure 4G). To confirm these observations, we re-analyzed these data by generating a mean consumption value for each subject within each block. Our evaluation of mean 2-hour ethanol consumption revealed a significant block*diet interaction [F(1,17)=62.85; p<.0001], which post-hoc tests confirmed was due to both within and between subjects differences at every level (Figure 5A). In contrast, although our evaluation of the first 15-minute period during DID revealed a significant block*diet interaction [F(1,18)=20.56; p=0.0003], post-hoc tests only identified differences within the first 2-week block (D1-D15; Figure 5B). Together, these findings suggest that something about the TL20 diet prevents animals from developing ethanol front-loading, but that once front-loading is established under the LD01 diet, it will persist. These data support our long-standing hypothesis that increases in the rate of ethanol consumption (i.e. front-loading) reflects an increase in motivation to experience ethanol’s post-absorptive effects and suggest that particular dietary attributes may increase or decrease ones susceptibility to developing an AUD.

## Discussion

Overall, these studies find that B6 mice maintained on LabDiet 5001 (LD01) reliably consume more ethanol, consume ethanol more quickly, and achieve higher BECs than mice maintained on Teklad 2920x (TL20). Furthermore, although differences in total fluid consumption were non-specific to ethanol (they were also observed in water-consuming groups), ethanol front-loading behavior was uniquely impacted in ways that support it as a learned behavioral characteristic related to experiencing ethanol’s post-absorptive effects. These findings provide additional evidence that rodent diet formulations can vastly impact voluntary ethanol consumption, and again highlight temporal drinking patterns as key to understanding excessive ethanol consumption and the development of AUDs. Overall, our work adds to the growing body of literature that laboratory rodent chow is anything but “standard” and should be reported in detail in all studies. There are, however, some important points that need consideration to bring the current findings into the broader context of the existing literature.

Given the non-specific effects of rodent diet on total fluid consumption using DID methods, as well as no observable differences in ethanol metabolism, it is not immediately clear what or how specific dietary factors might have contributed to differences in ethanol consumption. Even so, the data within this report are in close alignment with recently published work. Quadir and colleagues demonstrated that mice maintained on the LabDiet 5001 (LD01) consistently drank more ethanol than mice maintained on 3 other comparison diets using an intermittent access 24 hour 2-bottle choice paradigm (Quadir et al., 2020). Interestingly, they did not observe consistently greater *total* fluid consumption in the LD01 group, which our water group observations suggest might also be impacted. Another study substantiating diet’s impact on binge ethanol consumption assessed six different rodent diets and observed that animals consuming Teklad 2920 (TL20) routinely consumed the lowest levels of ethanol and had the lowest BECs compared with animals on the other five comparison diets (Marshall et al., 2015). Notably, LabDiet 5001 was not included in their study, and although there were no significant differences in water consumption among the 6 diets, it was lowest in mice maintained on TL20.

The specific attributes of the 2 diets under study that led to differences we observed remain unknown, but we considered several key factors. One we recognized early was that LD01 pellets are compressed whereas TL20 pellets are extruded. Extruded dietary components are typically ground up finer resulting in a less dense pellet (Kurtz & Feeney, 2020), which we observed also occurs in these two diets specifically, and is likely responsible for differences in size, shape and color of fecal boli between the different diet groups (Supplementary Figure 1). These density differences could potentially lead to differences in fluid adsorption or other physical barrier to rapid fluid ingestion. However, mice consuming TL20 were demonstrably capable of front-loading, they just only did so after this phenotype was established during maintenance on LD01 (Figure 4F).

We also carefully considered differences in ingredients. For example, the carbohydrate/protein ratio of food has been shown to impact ethanol consumption; specifically, that high carbohydrate to low protein ratio depresses ethanol consumption and vice versa (Kampov-Polevoy et al., 1999). Our observations are in-line with these findings, as TL20 has a higher carbohydrate/protein ratio than LD01, but these differences were extremely small and therefore unlikely to be a major driver of our observations. Despite similar proportions of macronutrients (Supplementary Table 1 and 2), the specific ingredients that were used in the manufacture of the two diets did vary quite substantially. Primary differences were the sources of protein between diets (Supplementary Table 1 and 2), with only LD01 containing soy-based products that are known to contain phytoestrogens (PEs). PEs functionally and structurally mimic mammalian estrogens and their active metabolites, which in turn can modulate estrogen-sensitive pathways (Mäkelä et al., 1995). In one study, animals fed a diet lacking PEs experienced deficits in learning and memory, which was rescued when subjects were supplied exogenous equol (a metabolite of estrogen; (Çalışkan et al., 2019). In contrast, animals fed a diet high in PEs experienced profound alterations in energy balance including reduced body weight, adiposity, and increased lipid oxidation (Cederroth et al., 2007). Our data potentially align with this report, as our subjects gained slightly more weight over a two-week period on the non-phytoestrogen-containing diet (TL20), as compared with the phytoestrogen high diet (LD01) (Supplementary Figure 2C). A recent mini-review on this subject was just published in which the authors suggested the existence of a direct relationship between phytoestrogens and ethanol consumption (Eduardo and Abrahao, 2021).

Relatedly, different commercial rodent diets have been found to lead to profound differences in the gut microbial community (Tuck et al., 2020). These diet-induced differences in microbial communities may impact the way food and other substances, like ethanol, are metabolized. Notably, BEC is influenced by how quickly ethanol is emptied from the stomach and the extent of metabolism that occurs after it passes through the stomach to the liver (Reviewed in (Zakhari, 2006). Since we did not thoroughly assess ethanol pharmacokinetics, diet-induced alterations in ethanol pharmacokinetics remain a possible mechanism of the behavioral differences. However, given we did not detect differences in ethanol metabolism, and we observed a significant positive linear relationship between ethanol consumption and BEC regardless of diet, we think this unlikely.

To our knowledge this is the first report that rodent diet can impact ethanol front-loading, and that these diet-induced alterations in front-loading may be primary to decreases in total ethanol consumption within a drinking session. Although additional work is necessary to describe the mechanisms of these observations, these studies provide further support of our long-standing hypothesis that increases in the rate of ethanol consumption (as indexed by front-loading) may reflect an increase in motivation to experience ethanol’s positive post-absorptive effects, and suggest that particular dietary attributes may influence ones susceptibility to developing deleterious patterns of ethanol consumption.

## References

Ardinger CE, Grahame NJ, Lapish CC, Linsenbardt DN (2020) High Alcohol–Preferring Mice Show Reaction to Loss of Ethanol Reward Following Repeated Binge Drinking. Alcohol Clin Exp Res 44:1717–1727.

Barbosa C, Cowell AJ, Dowd WN (2021) Alcohol Consumption in Response to the COVID-19 Pandemic in the United States. J Addict Med 15:341–344.

Çalişkan G, Raza SA, Demiray YE, Kul E, Sandhu KV, Stork O (2019) Depletion of dietary phytoestrogens reduces hippocampal plasticity and contextual fear memory stability in adult male mouse. Nutr Neurosci 1–12.

Carpenter RW, Padovano HT, Emery NN, Miranda R (2019) Rate of alcohol consumption in the daily life of adolescents and emerging adults. Psychopharmacology 236:3111–3124.

Cederroth CR, Vinciguerra M, Kühne F, Madani R, Doerge DR, Visser TJ, Foti M, Rohner-Jeanrenaud F, Vassalli J-D, Nef S (2007) A Phytoestrogen-Rich Diet Increases Energy Expenditure and Decreases Adiposity in Mice. Environ Health Persp 115:1467–1473.

Dole VP, Ho A, Gentry RT (1985) Toward an analogue of alcoholism in mice: criteria for recognition of pharmacologically motivated drinking. Proc National Acad Sci 82:3469–3471.

Eduardo PMC, Abrahao KP (2021) Food composition can influence how much alcohol your animal model drinks: A mini-review about the role of isoflavones. Alcohol Clin Exp Res.

Kampov-Polevoy AB, Garbutt JC, Janowsky DS (1999) ASSOCIATION BETWEEN PREFERENCE FOR SWEETS AND EXCESSIVE ALCOHOL INTAKE: A REVIEW OF ANIMAL AND HUMAN STUDIES. Alcohol Alcoholism 34:386–395.

Linsenbardt DN, Boehm SL (2015) Relative Fluid Novelty Differentially Alters the Time Course of Limited-Access Ethanol and Water Intake in Selectively Bred High-Alcohol-Preferring Mice. Alcohol Clin Exp Res 39:621–630.

Linsenbardt DN, Boehm SL (2014) Alterations in the rate of binge ethanol consumption: implications for preclinical studies in mice. Addict Biol 19:812–825.

Linsenbardt DN, Boehm SL (2012) Role of Novelty and Ethanol History in Locomotor Stimulation Induced by Binge-Like Ethanol Intake. Alcohol Clin Exp Res 36:887–894.

Linsenbardt DN, Boehm SL (2009) Agonism of the endocannabinoid system modulates binge-like alcohol intake in male C57BL/6J mice: involvement of the posterior ventral tegmental area. Neuroscience 164:424–434.

Linsenbardt DN, Moore EM, Griffin KD, Gigante ED, 2nd SLB (2011) Tolerance to Ethanol’s Ataxic Effects and Alterations in Ethanol-Induced Locomotion Following Repeated Binge-Like Ethanol Intake Using the DID Model. Alcohol Clin Exp Res 35:1246–1255.

Mäkelä S, Santti R, Salo L, McLachlan JA (1995) Phytoestrogens are partial estrogen agonists in the adult male mouse. Environ Health Persp 103:123–127.

Marshall SA, Rinker JA, Harrison LK, Fletcher CA, Herfel TM, Thiele TE (2015) Assessment of the Effects of 6 Standard Rodent Diets on Binge-Like and Voluntary Ethanol Consumption in Male C57BL/6J Mice. Alcohol Clin Exp Res 39:1406–1416.

Pollard MS, Tucker JS, Green HD (2020) Changes in Adult Alcohol Use and Consequences During the COVID-19 Pandemic in the US. Jama Netw Open 3:e2022942.

Quadir SG, Rohl CD, Zeabi A, Moore CF, Cottone P, Sabino V (2020) Effect of different standard rodent diets on ethanol intake and associated allodynia in male mice. Alcohol 87:17–23.

Tuck CJ, Palma GD, Takami K, Brant B, Caminero A, Reed DE, Muir JG, Gibson PR, Winterborn A, Verdu EF, Bercik P, Vanner S (2020) Nutritional profile of rodent diets impacts experimental reproducibility in microbiome preclinical research. Sci Rep-uk 10:17784.

(US) NAP (Ed.) (2003) Guidelines for the Care and Use of Mammals in Neuroscience and Behavioral Research.

Weerakoon S, Jetelina K, Knell G (2020) Longer time spent at home during COVID-19 pandemic is associated with binge drinking among US adults. Am J Drug Alcohol Abus 47:98–106.

Zakhari S (2006) Overview: how is alcohol metabolized by the body? Alcohol Res Heal J National Inst Alcohol Abus Alcohol 29:245–54.

