## Supplemental_Materials for "The Development, but not Expression, of Alcohol Front-loading in C57BL/6J Mice Maintained on LabDiet 5001 is Abolished by Maintenance on Teklad 2920x Rodent Diet"

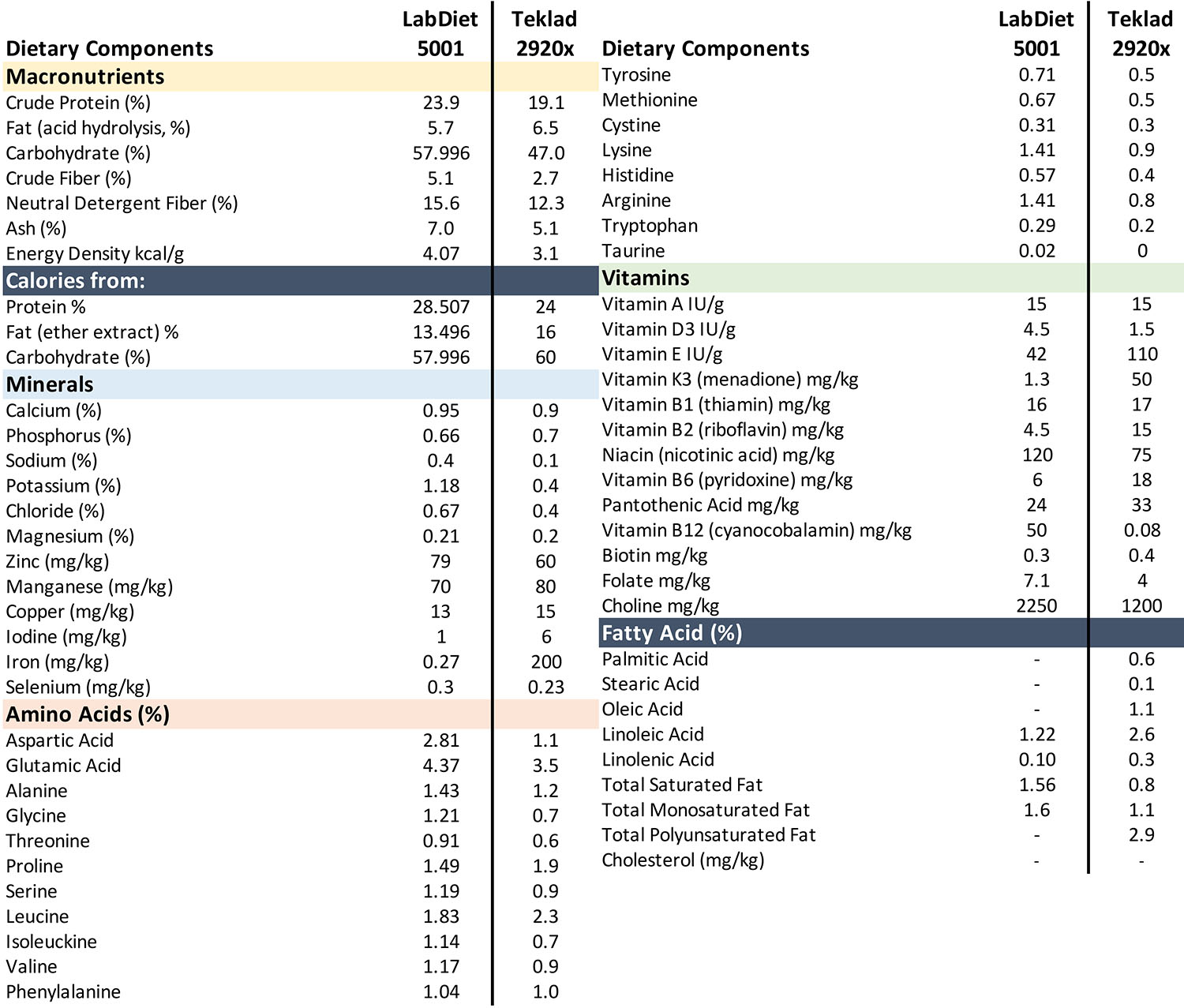

**Supplementary Table 1.** Detailed dietary components of two lab diets, Labdiet 5001 (LD01), and Teklad 2920 (TL20), all values listed in mg/kg of diet, percent, or units, as indicated.

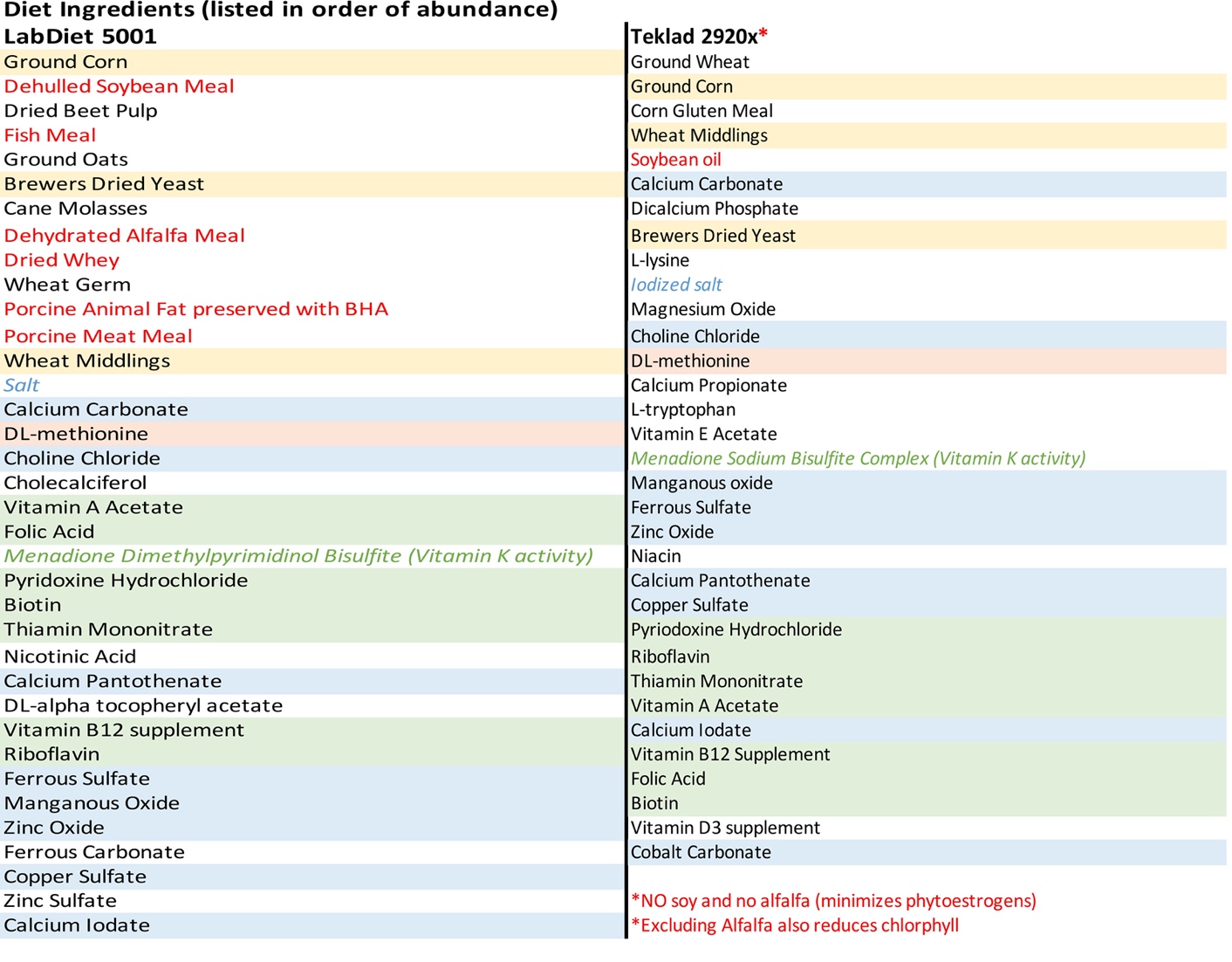

**Supplementary Table 2.** Detailed ingredient list of each diet, listed in order of abundance for LabDiet 5001 (LD01) and Teklad 2920 (TL20).

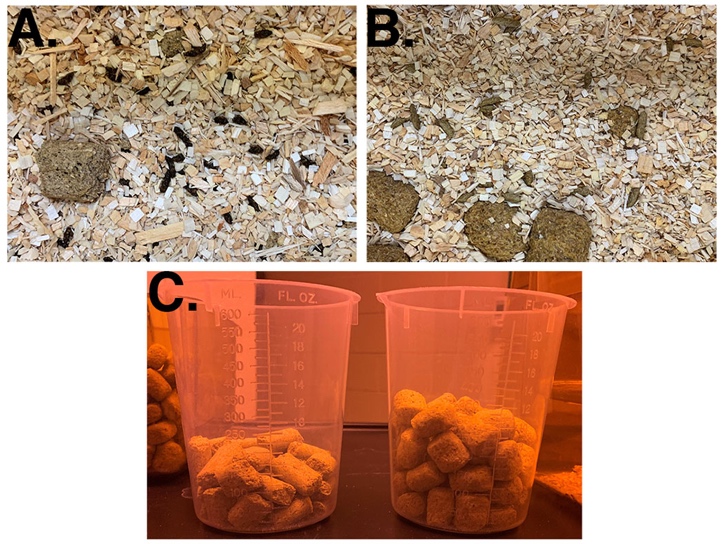

**Supplementary Figure 1**. Two different rodent diets influence fecal boli and have strikingly different densities. Fecal boli from LabDiet5001 (LD01) (**A**) appear black and small in appearance, whereas fecal boli from Teklad 2920x (TL20) (**B**) appear brownish and larger in appearance. Densities between the diets are also strikingly different, approximately 120 g of diet in left (LD01 vs right (TL20) in **C**.

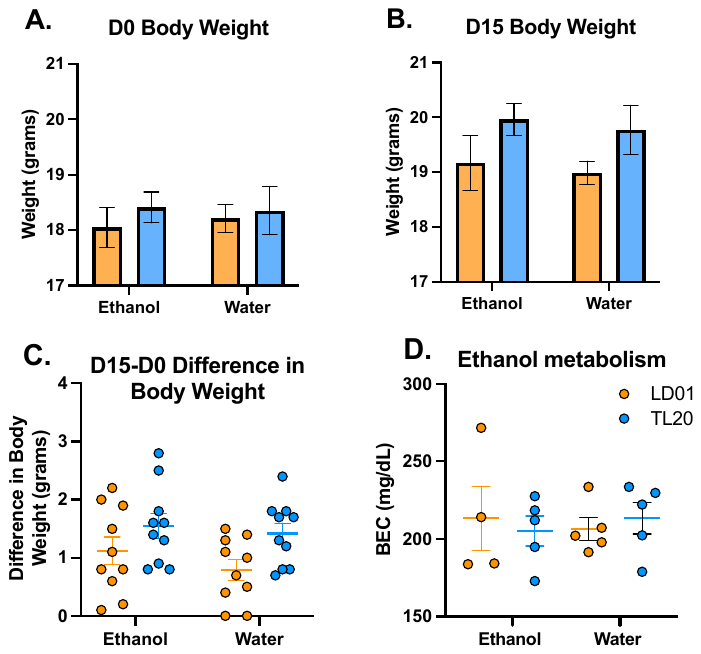

**Supplementary Figure 2.** Subtle but non-significant differences in female C57BL/6J mice maintained on either Labdiet 5001 (LD01) or Teklad 2920x (TL20) for a period of 2 weeks. Animal weights were taken at the beginning (D0) (**A**) and at the end of experiment 2 (D15) (**B**) were taken and revealed subtle but not substantial differences in weight gain (**C**). Maintenance on diet, nor previous fluid assignment affected ethanol metabolism (**D**).
